# Melodic expectation as an elicitor of music-evoked chills

**DOI:** 10.1101/2024.10.02.616280

**Authors:** Rémi de Fleurian, Ana Clemente, Emmanouil Benetos, Marcus T. Pearce

## Abstract

Music-evoked chills are pleasurable shivers, goosebumps or tingling sensations experienced during musical listening. Research on the characteristics of music perception that elicit chills has focused on low-level acoustic features in small samples of music. We collected a dataset of 1,019 pieces of music timestamped with onset times of 1,806 chills from 402 participants and used computational methods to predict chills onsets from simulations of low-level acoustic and high-level expectation-based elicitors. A machine learning classifier was trained to discriminate passages containing chills from those not containing chills. The results show that chills onsets are predicted better than chance and corroborate existing evidence for acoustic elicitors of chills with a much larger dataset. They also provide empirical evidence that music-evoked chills are elicited by surprise and uncertainty resulting from a high-level psychological process of expectation, which proved a more effective predictor than acoustic elicitors.

## 1 Introduction

Chills are shivers, goosebumps or tingling sensations that occur during certain kinds of sensory experience such as film, theatre, poetry, visual art, natural beauty as well as experience of nostalgia or inspiration [1]. Music-evoked chills (MECs) are chills experienced while listening to music. Evidence suggests that they are experienced by most individuals and are usually reported as being intensely pleasurable, often corresponding to moments of peak affective intensity while listening to a piece of music [2]. MECs have attracted scientific interest as an observable physiological response reflecting an unobservable internal state of pleasure while listening to music. MECs have proven to be robust activators of the brain’s reward system in neuroimaging experiments [3–5], neuropsychological comparisons [6, 7] and pharmacological interventions [8, 9]. However, the causes of MECs have remained challenging to understand.

A recent systematic review of research on MECs highlights a complex network of different influencing factors and different psychological mechanisms involved [2]. MECs are thought to be influenced by the interactive effects of three factors: the listener, showing particular demographics, traits and state [10, 11] which may be related to underlying genetic [12] and neural characteristics [8, 13]; the listening context, including social factors and concurrent tasks [14, 15]; and the music, containing particular characteristics whose perception can elicit MECs. Considering the last of these factors, specific elicitors related to psychological processing of the musical stimulus have been categorised into three classes [2]: low-level acoustic elicitors (e.g., perception of loudness changes), high-level musical elicitors (e.g., perception of structural change or development in melody, harmony, rhythm or dynamics) and emotional elicitors (i.e., perception of emotions expressed by music). These elicitors represent psychological processing of different cues in music with acoustic elicitors derived at a lower-level of auditory processing, reflecting perception of simple local changes in acoustic properties occurring at short timescales, while musical elicitors reflect higher-level perception of more complex auditory structures and relationships operating over longer timescales within a piece of music, which may be subject to effects of top-down processing (e.g., expectation, attention, musical experience) and the musical context. Emotional elicitors meanwhile involve recognition of the expressive emotional meaning communicated by a piece of music. Any of these elicitors may have an impact on emotional state, which in turn may influence occurrence of MECs [2], reflecting interactions between auditory perception and the limbic system [16].

These elicitors have been related to distinct psychological mechanisms involved in emotion induction by music [17–19]. Acoustic elicitors are related to brain-stem reflex mechanisms in which extreme or sudden changes in basic acoustic characteristics lead to rapid modulations of arousal, with MECs proposed to be elicited by peak levels of arousal [20, 21]. Musical elicitors have been associated with the psychological process of expectation during listening in which predictions are generated for forthcoming musical events (e.g., the pitch of the next note in a melody) [22, 23]. Emotional elicitors have been associated with the experience of being moved, a positive, empathetic sense of social bonding, involving communal sharing or closeness with others [24]. These mechanisms are not mutually exclusive and may coexist, either contributing in differing degrees to the same MEC episode or producing different kinds of MECs corresponding to each mechanism.

The present research aimed to understand the eliciting characteristics of music perception (i.e., psychological processing of the musical signal rather than individual differences between listeners or effects of context) that predict the onset of individual chills with a focus on acoustic and expectation-based elicitors (rather than emotional elicitors). Existing research reviewed below has produced evidence for these elicitors of MECs, which provides an empirical hypothesis-driven basis for selecting the computational simulations of acoustic and musical elicitors used in the present research.

Acoustic elicitors of MECs reflect perception of low-level properties of the auditory signal. Early research on MECs identified a relationship with sudden dynamic changes by analysing music scores for passages reported to elicit MECs by survey participants [25]. This was confirmed empirically in subsequent experiments [26–33] in which loudness was extracted from audio [or manually derived from music scores, 30] around the onset of MECs reported either retrospectively or continuously during listening. Loudness is by far the best-documented acoustic elicitor of MECs, but other acoustic elicitors have been identified using similar methods, suggesting that MECs correlate with increases in roughness, sensory dissonance or fluctuation strength [27–29, 32, 34], increases in sharpness or brightness [27–29, 31], high spectral centroid and spectral flux [27], high event density [27, 32, 33], or expansion of the frequency range in a high or low register [30, 33]. Using a causal intervention, Bannister [35] experimentally manipulated loudness and brightness in two musical passages that had elicited MECs in previous research [27]. In one of these passages, MECs were experienced more frequently if loudness was increased or brightness decreased, demonstrating a causal effect of loudness and brightness on MECs, as opposed to the correlational findings discussed above.

Musical elicitors of MECs reflect perception of high-level properties of musical structure. A seminal questionnaire study found that musical passages associated with MECs (in 57 pieces supplied by 83 respondents) included new or unprepared harmonies, sudden textural changes, melodic appoggiaturas, enharmonic changes, specific melodic or harmonic sequences, or prominent events arriving earlier than prepared for, in decreasing order of frequency [25]. These self-reported effects of melodic and harmonic features on MECs were confirmed in subsequent survey-based and empirical research [14, 15, 26, 27, 30, 36], notably through identification of effects of structural transitions and alterations such as changes in tonality [14]. Rhythmic properties [15, 37, 38] and vocals [14, 15] have also been implicated, although there is a lack of specificity regarding which exact properties were associated with MECs. Additional findings revealed effects of crescendi, build-ups and climaxes [14, 26, 27, 33, 37, 39], as well as textural changes [25, 26, 33, 37], notably through the entrance of new instruments or the interplay between solo and background instruments [14, 26, 27, 30, 36]. It has been argued that musical elicitors reflect a hypothesised effect of musical expectation on MECs [3, 17, 22, 23, 25, 40–43], since most of the musical elicitors of MECs reviewed above can be related to violations of expectation.

Expectation is the psychological process of generating implicit predictions for future events during listening based on the preceding musical context. Expectations can be generated for the pitch, timing or harmony of musical events. The psychological process of musical expectation is thought to reflect predictions derived from a learned internal representation of the syntactic structure of the music to which a listener is exposed [22, 44]. An effect of expectation on emotional and aesthetic experience of music has long been hypothesised [45, 46]. Empirical studies have demonstrated a relationship between expectation and affective states of arousal in musical listening [47, 48] while pleasure shows an inverted-U shaped relationship with expectation such that optimal pleasure is associated with intermediate levels of expectedness [49, 50]. Several theories have been proposed to account for this relationship between expectation and pleasure: optimal levels of arousal associated with tension and resolution [46, 51]; a balance between cognitive fluency arising from expectation confirmation, and contrastive valence arising from the limbic contrast between an automatic negative response to expectation violation and a subsequent neutral or positive response given the non-threatening nature of the stimulus [22]; positive reward prediction errors when the music is more rewarding than expected [23]; and optimal learning progress reflecting a balance between the reward associated with accurate prediction and the intrinsic reward of learning, associated with expectation violation [49, 52].

Emotional elicitors of MECs also deserve mention, including subjectively perceived valence, emotionality and meaning in music. Research has identified effects of emotional elicitors such as positive and negative valence [14, 39, 53, 54], perceived emotionality [27, 28, 39, 55] and perceived meaning [1, 14, 36]. However, such characteristics of musical stimuli are difficult to quantify precisely and objectively, especially as continuous features, since they rely on some degree of subjective interpretation and often relate to large sections of music rather than specific, identifiable events. For this reason, emotional elicitors were not considered in the present research, although they are likely to be potent elicitors of MECs and deserve further attention in future research. The Chills in Music (ChiM) dataset, accompanying a systematic review of research on MECs [2], consists of 1,022 musical pieces used in previous empirical research on MECs. These musical pieces were found to have more negative valence scores than a carefully matched set of tracks [56]. However, the ChiM dataset lacks precise timing of MEC onsets and consistent information about the exact versions of the pieces of music found to elicit MECs, so could not be used in the present research. Mori [38] asked 54 participants to listen to individually self-selected pop songs while indicating MECs or tears with button presses. Using multi-class ridge regression and a range of acoustic features, a better-than-chance classification accuracy of 43% was achieved for MEC onsets. Classification performance was driven most strongly by the minor mode immediately preceding MECs (highlighting a possible relationship with expressed valence [56]), higher event density and rhythmic entropy at the onset of MECs, and higher spectral flux and rhythmic entropy after the onset of MECs.

This review of existing research provides evidence for effects of specific acoustic elicitors on MECs but for relatively small samples of participants and musical stimuli. Research has also identified higher-level musical elicitors of MECs, many of which can be related to expectation. While expectation has long been hypothesised to be an important elicitor of MECs [25], this theoretical prediction has yet to be robustly tested experimentally. In the present work, we address both issues by creating a large dataset of 1,019 pieces of music supplied by 402 individuals containing precise onset times for 1,806 MECs, which we predict with machine learning models that combine computational simulations of low-level acoustic and high-level expectation-based perception hypothesised to elicit MECs based on the research discussed above. The results show effects of both low-level acoustic and high-level expectation-based elicitors, with expectation-based elicitors proving more effective predictors than acoustic elicitors.

## 2 Results

### 2.1 Features

Both acoustic and musical elicitors of MECs were simulated computationally. To simulate acoustic elicitors, low-level features were computed from the audio files for each track. The acoustic features included: loudness using a measure of the spectral envelope (Envelope); timbre using measures of roughness, flatness, brightness, spectral centroid, spectral flux, spectral entropy and spectral spread; tonality using measures of key distance and harmonic change (HCDF); and presence of vocals (Vocals). As described in the Introduction, the impact of these simulated acoustic elicitors on MECs was hypothesised based on previous research [25–35].

To simulate expectation as a musical elicitor, we used a computational model of auditory expectation [44] that has been the subject of substantial empirical evaluation and has proved capable of simulating expectation in music perception across a range of experiments [44, 47, 49, 50, 57–60]. The model simulates expectations as a psychological process of probabilistic prediction based on statistical learning of the structure of the music to which it is exposed (see Methods). Dynamic expectations derived from perception of repeated structure within a piece of music are simulated through online learning within each track while schematic expectations derived from long-term experience of a musical culture are simulated by training the model on a corpus. In the present research, we use a corpus of Western tonal folk songs and hymns which previous research has demonstrated produces a model that accurately simulates perception [47, 49, 58, 59] (see [57] for a review).

The trained model generates a conditional probability distribution for each note in a melody given the preceding melodic context. Given these distributions, the high-level expectation-based features used in the present research are melodic entropy and melodic information content (see Methods for further details). Melodic entropy reflects the model’s prospective uncertainty in making probabilistic predictions for the pitch of the next note in a melody. Information content (the negative log probability) reflects the unexpectedness or surprisal of the pitch that actually follows. As described in the Introduction, the impact of these simulated expectation-based elicitors on MECs was hypothesised based on previous research [22, 23, 25, 45, 46].

### 2.2 Feature differences between MEC and non-MEC excerpts

The stimuli were segmented into two unbalanced sets of 20-s excerpts: an MEC set capturing 10-s before and after each MEC and a control set capturing most other moments within the tracks. Figure 1 shows changes of each individual feature in these 20-s excerpts distinguishing MEC from control excerpts. Significant differences were obtained by comparing the observed differences with a null distribution computed by permuting the labels 5,000 times (see Methods). The results can be divided into four categories: features that were consistently higher than control excerpts around MEC onsets; features showing a sharp increase around MEC onsets; features showing a significant increase or decrease around an MEC onset followed by a return to the level preceding the MEC; and features that showed no significant differences from control excerpts.

**Figure 1:**
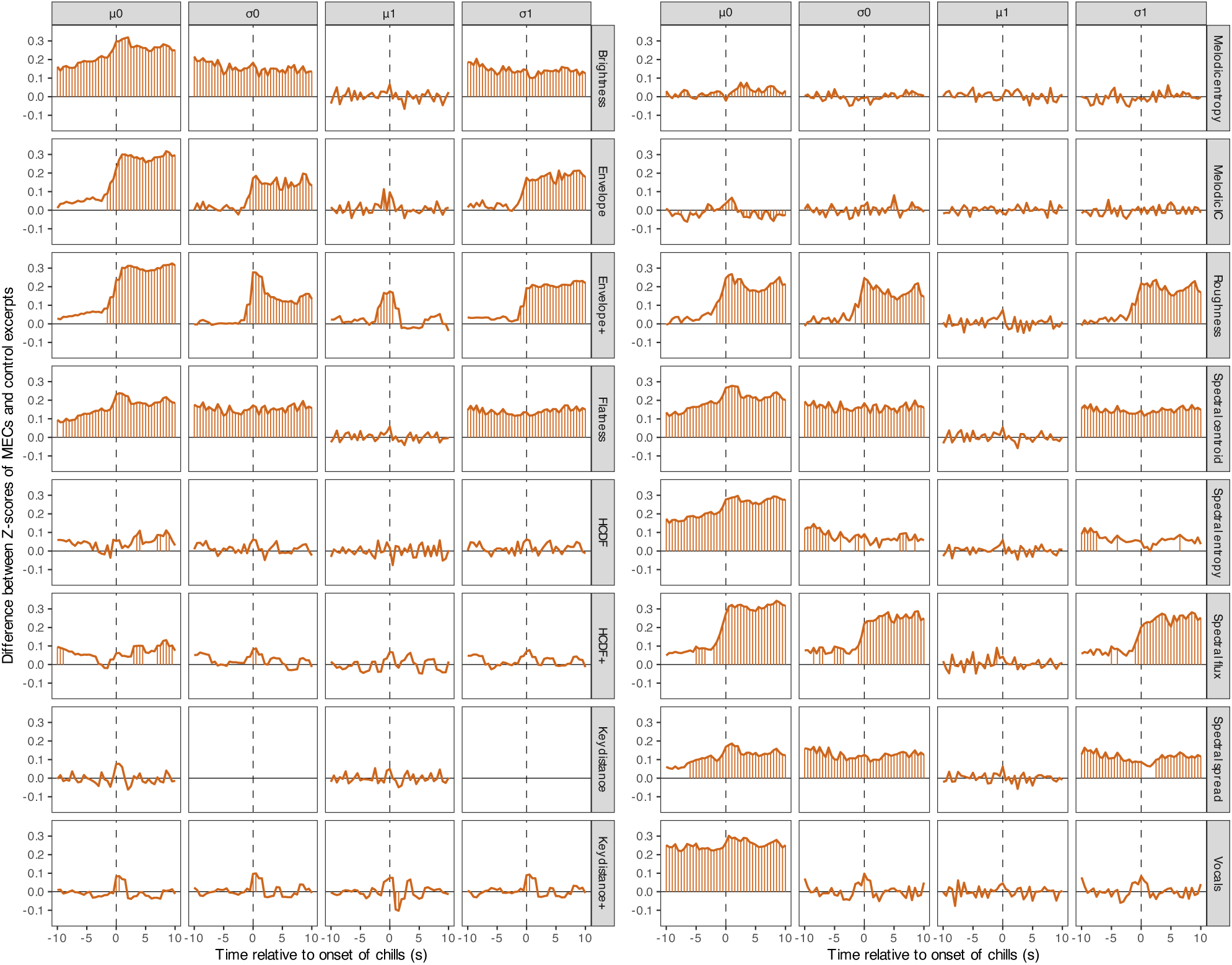
Each row of plots corresponds to features representing a simulated elicitor of MECs (see Methods), with each of the four columns corresponding to the mean (*µ*) and variability (*σ*) of the zeroth (0) and first derivative (1), respectively. The x-axes represent 20-s excerpts centred around MEC onsets (*x* = 0). The y-axes represent the difference between normalised feature values (Z-scores) averaged over all excerpts containing MECs and equivalent feature values averaged over all control excerpts. The horizontal line at *y* = 0 represents no difference in Z-score between MEC and control excerpts. Vertical lines denote frames for which the difference is significant according to frame-by-frame permutation tests. Permutation tests were not conducted for key distance variability due to the way these features were computed (see Methods). Source data are provided as a Source Data file.

Examining first the averaged features (identified by *µ*_0_), MECs were characterised by elevated brightness, flatness, spectral centroid, spectral entropy, melodic entropy and presence of vocals, as well as sharp increases in envelope, roughness, spectral flux and spectral spread. While several frames also showed significant differences regarding the harmonic change detection function (HCDF) and its associated feature computed with a larger sliding window (HCDF+), the overall differences were less convincing than for other features. Key distance (only when computed with a larger sliding window: Key distance+) and melodic information content significantly increased while melodic entropy significantly decreased around MEC onset followed by a return to their pre-MEC level, suggesting that MECs are elicited by isolated musical events with increased key distance and melodic information content but reduced entropy.

The variability in these features (identified by *σ*_0_) largely followed similar patterns. The only exception was the presence of vocals, which only contained a single frame with a significant difference between MECs and controls, located at the exact onset of MECs. Note that the presence of vocals was the only binary feature and relatively slow-moving compared to the other features due to how it was preprocessed. Nonetheless, this can be interpreted as vocals being more likely around MEC onsets (as seen with *µ*_0_) and MECs being slightly affected by increased variability in the presence of vocals around their onsets.

Significant effects were much more sparse when examining the rate of change of each feature (identified by *µ*_1_). A convincing peak only occurred for envelopes computed with a larger sliding window (Envelope+), providing further evidence that sudden, large changes in loudness are associated with MECs. Variability in these rates of change (identified by *σ*_1_) followed almost exactly the variability in the original features (*σ*_0_), consistent with the intuition that when variability in a feature substantially increases, this is also reflected in the variability of its rate of change.

### 2.3 Classifying MECs from acoustic and expectation-based elicitors

The dimensionality of the feature set (62 features as shown in Table 1) was reduced using PCA with 15 principal components retained (see Methods). These were used to train a linear support vector machine (SVM) to distinguish positive (MEC onsets) from negative (control) labels. Positive labels were associated with 500-ms frames containing a reported MEC onset as well as the two frames before and after this frame (see Methods and Supplementary Information). The SVM was trained and tested by splitting the data track-wise using five-fold cross-validation. Each track was randomly assigned to one of five folds with 203–204 tracks and 340–371 MEC occurrences per fold.

**Table 1:** List of features included in both sets of analyses.

| Tool | Feature category | Features |  |  |  |
| --- | --- | --- | --- | --- | --- |
| | | $\mu_0$ | $\sigma_0$ | $\mu_1$ | $\sigma_1$ |
| MiningSuite | Envelope | ✓ | ✓ | ✓ | ✓ |
|  | Envelope+ | ✓ | ✓ | ✓ | ✓ |
|  | Roughness | ✓ | ✓ | ✓ | ✓ |
|  | Flatness | ✓ | ✓ | ✓ | ✓ |
|  | Brightness | ✓ | ✓ | ✓ | ✓ |
|  | Spectral centroid | ✓ | ✓ | ✓ | ✓ |
|  | Spectral flux | ✓ | ✓ | ✓ | ✓ |
|  | Spectral entropy | ✓ | ✓ | ✓ | ✓ |
|  | Spectral spread | ✓ | ✓ | ✓ | ✓ |
|  | Key distance | ✓ |  | ✓ |  |
|  | Key distance+ | ✓ | ✓ | ✓ | ✓ |
|  | HCDF | ✓ | ✓ | ✓ | ✓ |
|  | HCDF+ | ✓ | ✓ | ✓ | ✓ |
| Spleeter | Vocals | ✓ | ✓ | ✓ | ✓ |
| IDyOM | Melodic entropy | ✓ | ✓ | ✓ | ✓ |
|  | Melodic IC | ✓ | ✓ | ✓ | ✓ |

Note. Four types of features for each feature category. $\mu_0$ = mean of the original values, $\sigma_0$ = standard deviation of the original values, $\mu_1$ = mean of the first-order difference, $\sigma_1$ = standard deviation of the first-order difference, + = feature computed using a longer, sliding window, IC = information content.

For the evaluation, each track was segmented into consecutive 5-s segments, labelled positive if they contained at least one 500-ms frame classified as positive by the SVM and negative otherwise, while the ground truth consisted of the same 5-s segments labelled positive if they contained an actual MEC onset and negative otherwise (see Methods). The area under the receiver-operating characteristic curve (AUC) is shown in Figure 2, reflecting classification performance for each fold, along with the threshold for the highest F*_β_* values for each curve using *β* = 2. In terms of overall performance, the SVM exhibited an AUC of 0.597, F*_β_* of 0.167, and balanced accuracy (the arithmetic mean of the accuracy in each class) of 0.58. See the Supplementary Information for a model comparison. One-sided permutation tests in which the trained models were re-evaluated on 1000 variations of the test set with randomly permuted labels show that AUC and balanced accuracy are significantly better than chance performance, p *<* .001.

**Figure 2:**
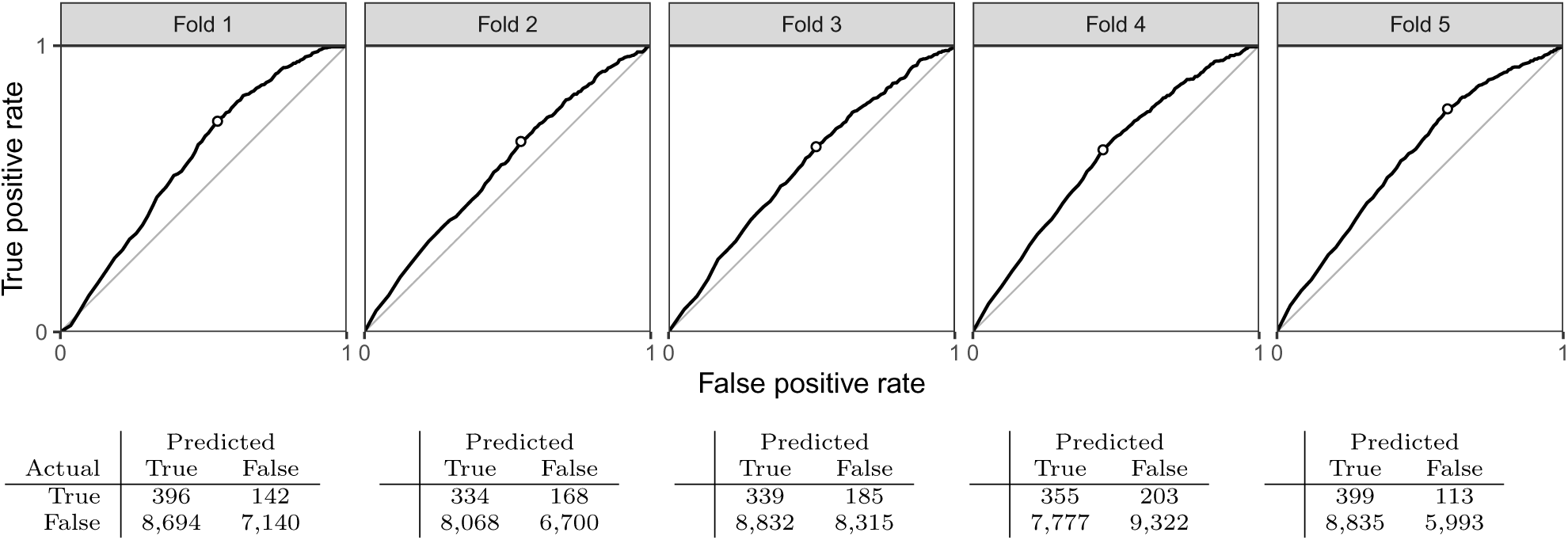
Receiver-operating characteristic (ROC) curves for each cross-validation fold of the trained SVM. Overall AUC for the model was computed by averaging the AUCs for each curve. The threshold returning the highest F*_β_* value is visualised by a circle on each curve and corresponds to the point at which all evaluation metrics were retained before being averaged to return overall evaluation metrics for the model. Chance performance is indicated by the line of equality (*x* = *y*), with curves above this line indicating that the true positive rate (recall, sensitivity, hit rate or power) exceeds the false positive rate (fall-out or type-I error). Across all tracks, 82,310 5-s segments were evaluated of which 2,634 were positive (mean per track = 2.58, sd = 1.77) and 79,676 negative (mean per track = 78.19, SD = 111.78). Under each figure is the confusion matrix for predicted and actual MEC segments, showing true positives, false negatives, false positives and true negatives (from left to right by row). Source data are provided as a Source Data file.

Finally, the contribution of each feature to the SVM in terms of feature importance (see Methods) is visualised in Figure 3. Melodic entropy and information content were the best predictors of MECs, with melodic entropy reaching a value of 1, meaning it was the most important predictor on all five cross-validation folds. Variability of the first-order difference of melodic entropy, mean spectral flatness, spread, and centroid followed these features. The least contributing features were mean first-order differences of brightness, roughness, spectral entropy, spread and centroid.

**Figure 3:**
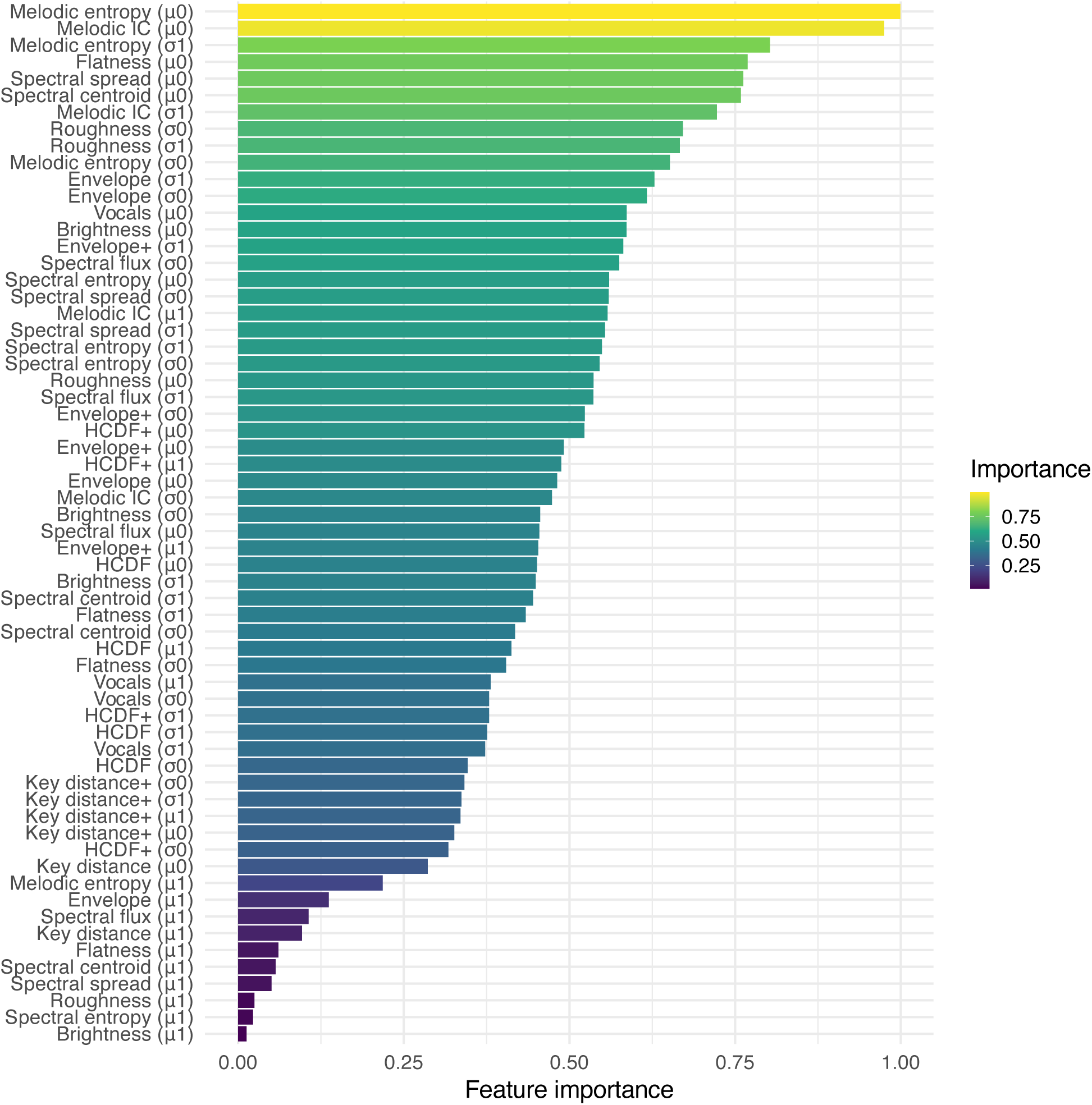
Relative feature importance for the SVM. Feature importance is rescaled linearly such that the feature contributing the most to each cross-validation fold receives a score of 1 (e.g., mean melodic entropy) and the feature contributing the least receives a score of 0, which no feature did for all five folds. Source data are provided as a Source Data file.

## 3 Discussion

### 3.1 Melodic expectation as an elicitor of MECs

These results provide convincing evidence that the stimulus-based perceptual elicitors of MECs include mechanisms related to pitch expectation, newly demonstrated here, alongside perception of acoustic cues, as demonstrated in previous research but corroborated here using machine learning methods on a much larger sample of participants and musical stimuli. In particular, expectation-based measures of pitch surprisal and predictive uncertainty increased classification performance (see Supplementary Information) and were convincingly higher in feature importance than acoustic features such as loudness, timbre (spectral flatness, centroid and spread), roughness and presence of vocals. The permutation testing suggests that MECs are elicited by surprising events (indicated by an increase in information content at MEC onset) in the context of relatively low surprisal (lower information content than baseline surrounding the MEC onset). MECs are also associated with generally higher melodic entropy (uncertainty) than non-MEC passages. This corroborates theoretical proposals dating back more than 150 years suggesting a prominent role of expectation in aesthetic experience of music [45, 46, 49, 50] and more recently in MECs specifically [17, 23, 25, 41, 43].

The permutation tests for individual acoustic elicitors replicated many of the findings from prior research. These results are not exploratory since we only included acoustic elicitors hypothesised from previous research. In addition, we applied strict Bonferroni correction, reducing the alpha threshold for statistical significance to slightly over 0.001, therefore providing a high degree of confidence in the identified effects. Tying these results back to findings from prior research [2], the permutation test analysis provided a large-scale replication of effects showing MECs as being associated with all the acoustic and expectation-based musical elicitors simulated by extracted features, including increases in loudness [25], crescendi [39], increased roughness [29], brightness [27], event density [32], spectral centroid and flux [27], expansion of the frequency range [30] as well as changes in texture, harmony and tonality [25]. The most important acoustic features were the mean level of spectral features (flatness, spread and centroid) along with variability in roughness and amplitude envelope. The results for harmonic change and key distance were less convincing than for other acoustic features, consistent with previous findings [38], perhaps due to their low-level nature. The results provide quantitative evidence that MECs are more likely in the presence of vocals. Other research has found no effect of the content of lyrics on MEC frequency [38], so it may be that the effect observed here is related to the acoustic or musical characteristics of the vocal line rather than lyrical content. On the other hand, survey respondents have identified lyrics (including narrative and emotional content) as a contributor to MECs [14, 15] suggesting that further research is warranted combining the present approach with measures of lyrical content.

The expectation-based measures of mean melodic entropy and information content were the best predictors of MEC onsets by far and the variability in their rates of change also achieved feature importance levels greater than all but three of the acoustic features. This indicates that MECs are associated with points in a melody that have high mean levels (*µ*_0_) of predictive uncertainty (entropy) and surprisal (information content). Furthermore, while mean change (*µ*_1_) in melodic IC/entropy within a 500-ms window has lower importance, the corresponding variability in this measure of change (*σ*_1_) achieved higher importance than simple variability (*σ*_0_). This suggests that variability in changes of IC/entropy are more important than variability of IC/entropy per se. However, since the two measures of variability (*σ*_0_ and *σ*_1_) are correlated (IC: r(544,914) = .57, p *<* .001, 95% CI = [.5679, .5715]; entropy: r(544,914) = .59, p *<* .001, 95% CI = [.5917 .5952]), these results are best interpreted as indicating that (some measure of) variability of IC/entropy is predictive of MECs in addition to the absolute level of IC/entropy.

The strong performance of expectation-based measures comes despite them being computed from melodies automatically extracted from the audio tracks—as manual transcription was not possible given the size of the dataset. Automatic melody transcription from audio is a notoriously challenging problem in music informatics [61] and the method used produced much less accurate input data than for the acoustic measures for which raw audio files were available. These inaccuracies are likely to make the melodies more unpredictable (increasing IC and entropy) but there is no reason to believe that this unpredictability would systematically co-vary with occurrence of MECs. Rather, it is likely that it adds noise to both MEC and non-MEC segments making it harder for the classifier to distinguish one from the other. Therefore, the effects of expectation found here are likely to represent a significant underestimate of the true effects. A possibility for future research would be to run a smaller-scale study using manual transcriptions of a subset of the oChim dataset and compare them to the automated transcriptions to quantify the noise due to the transcription method and its impact on the relationship between expectation and MECs. Furthermore, the inaccuracy of melody extraction precluded examining expectation-based measures for rhythm, an interesting avenue to follow in future research, along with harmonic expectation [48, 50]. Another potential source of inaccuracy is the corpus used to train the computational model of expectation; although this has been shown to simulate the expectations of Western listeners in previous research [44, 47, 49, 57–59], it does not represent exactly the music that each participant has listened to through their lifetime, informing their melodic expectations. While research has found that models trained on particular styles better simulate expectations of listeners with experience in those styles [59, 60], an interesting direction for future research is to examine training corpora that reflect specific individual listening histories. It may even be that effects of expectation work in different ways for the musical styles represented in the present work (e.g., rock compared with classical) such that different stylistic training corpora would produce more accurate predictions for pieces in different styles. Finally, future work might also benefit from extracting beat-aligned acoustic features as opposed to frame-based representation in the time domain; this could help integrate low-level acoustic elicitors with high-level expectation-based processing.

Multiple factors are thought to influence MECs, including emotional state [21, 24], individual differences between participants [10, 11] and the listening context [14, 15], all of which could have influenced the MECs reported here. Against this backdrop, which may have added considerable noise to the analysis, significant effects of acoustic and expectation-based predictors on MECs are all the more notable. The use of PCA combined with the fact that melodic IC and entropy were not correlated with any other features (see Figure 7) will have addressed collinearities between features to some extent. However, the effect of expectation-based elicitors may still depend in some way on the presence of acoustic elicitors since these were not removed or controlled in the music. Furthermore, it is possible that the effects of acoustic and expectation-based elicitors depend to some extent on their impact on emotional experience which in turn influences MECs. Although the present results shed light on the relative importance of acoustic and expectation-based elicitors of MECs, this does not extend to the relative importance of these elicitors with other factors contributing to MECs, including emotional elicitors. In future research, this could be investigated in an experiment that added continuous measurements of perceived emotion expressed by music to the design used here.

### 3.2 Statistical analyses

The fact that the SVM classifier performed above chance provides convincing evidence that both acoustic and expectation-based processing contribute to eliciting MECs. However, while recall was high, precision was low for all models. Precision refers to the proportion of classified MECs actually annotated as MECs in the dataset, while recall refers to the proportion of MECs in the dataset classified as MECs. In other words, the trained model identified a large proportion of the MECs in the dataset (high recall) but also predicted too many MECs at points where none were recorded in the dataset (low precision), although all the models still performed better than chance as shown by all AUC values exceeding 0.5. It is because of this low precision that we opted to report results based on the highest F*_β_* value returned by each model, since not doing so disproportionately penalised recall by optimising for small increases in precision. Optimising for recall also made sense conceptually due to the nature of the data. While we can be sure that each MEC recorded represents an experience of chills for at least one person, we cannot infer from the absence of an MEC report that chills would never be experienced by anyone at that point in a particular piece of music. If we were to massively increase the sample size, it seems likely that additional onsets of MECs would have been reported for some of the passages used as controls for model training. This makes the ground truth incomplete, which motivated attempting to predict all the MECs reported in the data by maximising recall (rather than predicting only the MECs present in the data by maximising precision), with the added advantage that an F*_β_*-optimised model should provide more useful information about which features best characterised MEC onsets when looking at feature importance.

Feature importance provides interesting insights when combined with the results of the permutation tests, by allowing comparisons between the magnitude of the difference between MECs and controls in each feature, and the degree to which these differences were predictive of MECs. For instance, the lack of detected effects of first-order differences on MECs in the permutation testing is also apparent in the fact that most of these features were not strong predictors of MECs in the classification analysis. However, in some cases, these comparisons are less easy to interpret. Many features (e.g., sharp increases in loudness) benefit from extensive prior empirical support and strongly displayed the hypothesised behaviour in the permutation tests, but they were not highly ranked in terms of feature importance in the SVM analysis. Conversely, the magnitude of the effects detected in the permutation tests for melodic entropy and information content was much lower than for other features, and yet, these expectation-related features were the best predictors of MECs. This suggests that, while previous correlational evidence for an association between some acoustic features and MECs was replicated, these features were not always strong predictors of MECs.

There could be several reasons for these differences between the permutation test analysis and the SVM classification analysis. First, the two analyses used different analysis windows centred around each MEC: 20 s for the permutation tests computed on individual frames and 2 s for the SVM classification, meaning that effects in the permutation tests falling outside a 2-s window would not have been considered by the SVM, which takes account of all frames within the 2-s window. These analysis windows reflected the different motivation and goals for each analysis: examining temporal evolution of feature prediction for the permutation test and discrimination of MECs from non-MECs based on a local window for the SVM analysis. The SVM classification also differs from the permutation tests in that it is multivariate rather than univariate, meaning that it could be affected by collinearity between features. Although PCA was used to avoid such effects, it is possible that the SVM was affected by residual relationships between principal components. A more likely possibility is that the differences reflect divergence in the analysis methods. Taking loudness as a concrete example, it seems likely that increases and decreases in loudness are both (approximately equally) common in music but that loudness increases are more common around MECs. This would mean that in the permutation tests, the average loudness for MEC segments would be greater than that for non-MEC segments, which would average to approximately zero given the roughly equal number of both increases and decreases in loudness. By contrast, the SVM is trying to classify episodes as MEC or non-MEC on a case-by-case basis without averaging, so the common occurrence of loudness increases would mean that they are not actually predictive of MEC onsets due to the large number of false positive instances (i.e., loudness increases that are not associated with an MEC). Conversely, small differences in expectation between MECs and controls could be due to small (but consistent) effect sizes or to increases in entropy and information content only being present in a subset of MEC onsets. Nonetheless, these expectation-related elicitors appear to be highly specific to MECs, given their feature importance in predicting chills onsets.

### 3.3 Neural basis

MECs have been shown to be associated with activation of the reward system of the brain including the ventral striatum [3, 4, 13], insula [4, 13] and amygdala [4, 13]. The experience of pleasure during MECs has been specifically linked to dopaminergic activation in the ventral striatum using a combination of ligand-specific PET and fMRI while activation in the dorsal striatum coincides with anticipation of MECs [3]. Furthermore, pharmacological manipulation of dopaminergic function with a dopamine precursor (levodopa) and antagonist (risperidone) respectively increase and decrease MECs [8]. A recent study showed that duration of MECs was predicted by right-hemisphere resting state functional connectivity between auditory cortex and the reward system (nucleus accumbens, orbitofrontal cortex and amygdala) in the 40 s prior to the start of the music [16]. From a theoretical perspective, it has been proposed that expectation plays an important role throughout the auditory brain, connecting low- and high-level perception of pitch and timing respectively within the ventral and dorsal auditory cortical pathways [62, 63], as well as their interactions with the reward system [64] associated with chills [23]. Specifically, sensory expectations for musical events of the kind simulated here generate prediction errors in auditory pathways which in turn generate positively or negatively valenced reward prediction errors that modulate states of pleasure associated with chills [23]. Based on this theoretical framework, we hypothesise that the acoustic and expectation-based elicitors investigated here correspond to neural processing in earlier and later stages respectively of the ventral auditory pathway from primary auditory cortex to inferior frontal gyrus.

### 3.4 Limitations

This study is not without limitations, foremost of which is the generally poor classification performance of the SVM. However, it is worth placing model performance within the context of the task at hand. Automatic detection of MEC onsets is far from trivial, due to the inherent impossibility of collecting exhaustive data and MECs being a subjective, psychophysiological response known to be driven by a wide range of elicitors, some of which would be exceedingly difficult to quantify for modelling purposes. For example, there is evidence that MECs are influenced by the personality and current affective state of the listener [65] and their context [14, 66, 67]. There is also evidence that MECs are influenced by emotional experience [2] for which expectation is just one of several mechanisms by which music can induce an emotional state [17]. Another factor is the potential influence of individual episodic memories for particular pieces of music. Following Salimpoor et al. [68], we attempted to limit the effect of personal memory in the instructions to participants but this may not have been entirely effective. Given the complexity of these contributors to experience of MECs, it is striking that we find significant effects of stimulus-driven acoustic and expectation-based elicitors.

Future research should aim to understand better how the acoustic and expectation-based elicitors of chills investigated here combine with emotional elicitors and episodic memory as well as individual (personality, demography, emotional state) and contextual factors (e.g., social context). For example, the individual trait of reward responsiveness is related to the intensity of MECs [11] while a recent study investigated in a large sample of 2,937 individuals whether occurrence of chills during presentation of 20 audio and 20 audio-visual stimuli (some of which included music) could be predicted by emotional state, demographic factors and a wide range of personality traits [10]. The results showed that chills were associated with greater arousal and valence as well as greater stimulus familiarity but also personality traits (greater extroversion, conscientiousness and disposition towards positive emotions) and demographic factors. Since that study did not record the particular time points at which chills occurred, an interesting direction for future research would be to investigate how these factors combine and interact with the acoustic and expectation-based elicitors of MECs investigated in the present research. In this context, it would also be worthwhile replicating this study with a broader range of musical cultures, given the preponderance of European and North American participants (87%) and music in the present work.

Another limitation arises from the fact that we wanted our analyses to be hypothesis-driven and interpretable, which meant that we needed to use both interpretable features and interpretable models at the necessary cost of predictive performance. It is highly likely that more powerful approaches (e.g., neural networks, ensemble machine learning methods) or features such as mel-frequency cepstral coefficients [38] would result in better predictive performance. However, such methods would have significantly compromised the interpretable hypothesis-driven nature of the research. Future work will gain from methodological insights gathered in the present work (see Methods and Supplementary Information), such as the use of features computed over large sliding windows and segmented using a 500-ms frame size, or the use of segment-based metrics for evaluation. Exploring the benefits of using larger frame sizes might also lead to improved performance. It may turn out that there is a relatively low ceiling to predictive performance based on acoustic and expectation-based musical elicitors. If so, this would suggest a more important role for emotional elicitors than previously anticipated. Considering the extent of previous findings about the relationships between MECs and various extra-musical factors such as personal meaning or the state of being moved [2], it is possible that only a small proportion of MECs are exclusively driven by acoustic and musical elicitors via the psychological mechanisms of brain stem reflex and musical expectation [17]. Future work should attempt to add emotional elicitors to the analysis of acoustic and musical elicitors as reported here.

Finally, the present work relies on computational modelling to simulate acoustic and expectation-based elicitors of MECs. While the computational model used to simulate melodic expectation has been extensively tested in empirical research [44], a question remains about the perceptual validity of the audio features extracted to simulate acoustic elicitors. It is common in psychology experiments to extract audio features using existing music information retrieval toolboxes–––as a rough example, MIRtoolbox [69] has been cited in more than 600 articles including the word *psychology*. However, it is worth noting that many of these features were designed and evaluated for practical tasks, and, therefore, might not necessarily equate to human perceptual representations of sound [70].

In summary, the present work yields evidence that both low-level acoustic and high-level expectation-based elicitors influence the occurrence of music-evoked chills. In contrast to previous research, large samples of participants and pieces of music were collected with precise time stamps for MECs, and machine learning methods were used to investigate whether computational simulations of acoustic and expectation-based elicitors were predictive of MEC onsets. The results both replicate previous findings for acoustic elicitors of MECs and provide evidence for the role of high-level musical expectations, demonstrating that simulated surprise and uncertainty distinguish MECs from other musical passages and highlighting the greater importance of expectation-based elicitors relative to acoustic elicitors in classification performance. Expectation mechanisms are of broader interest beyond music perception since expectation is thought to be an important general component in neural computation [71], operating in other perceptual domains such as language [72, 73] and visual perception [74, 75]. This means that expectation may generalise as an elicitor of chills in other perceptual domains (e.g., visual perception of dance or reading of poetry) to a greater extent than acoustic elicitors, which are by their nature specific to auditory processing. Future research should seek to explore the many remaining gaps in knowledge about the relationship between expectation and MECs. Notably, questions remain about the exact interaction between uncertainty and surprise, the role of harmonic and rhythmic expectation, the differences between schematic and veridical expectation, and the interactions between acoustic and musical elicitors, stylistic preference and familiarity [4, 27, 29, 35, 65], and affect [2].

## 4 Methods

### 4.1 Dataset

The *Onsets of Chills in Music (oChiM)* dataset accompanying the present study was collected via an online survey to access a broad sample of individuals and music. Having provided informed consent, participants reported basic demographic information such as age and country of residence. Given the lack of any convincing theoretical basis or empirical evidence for an effect of sex or gender on music-evoked chills, these were not considered in the study design [2]. Participants were then asked to provide up to ten pieces of music that often evoke chills (defined as shivers, goosebumps or a tingling sensation experienced in response to music listening). For each piece of music, participants provided the name of the artist or composer, the title of the piece of music, a URL for the piece of music (streamed on YouTube, Spotify, SoundCloud, or a similar platform), and between one and five precise timestamps for the instants at which they frequently experience the MEC onset (i.e., the exact moment at which chills begin). Audio files were retrieved from these URLs and used to compute the predictors of MECs used in the analysis (see Methods: Feature Extraction). The oChiM dataset contains the URL and the extracted predictors but not the audio files themselves.

The focus of the present experiment is the role of the stimulus and its perception in eliciting MECs but chills are also thought to be affected by episodic memories of past experiences of particular pieces of music [1]. As an experimental measure to factor out such effects, participants were advised not to include music that could be strongly associated with specific personal memories such as life events (e.g., a wedding, a memorable concert) or periods of time (e.g., last summer, secondary school), or music that was taken from a film soundtrack if they had watched the film in question. These restrictions (taken from a previous study by Salimpoor *et al.* [68]) were intended to maximise the likelihood that the MECs were induced by detectable acoustic and musical elicitors, with the potential to generalise across individual participants, as opposed to individual autobiographical elicitors driven by episodic memory, which are unlikely to generalise across individuals.

The survey was hosted on Qualtrics (Qualtrics, Provo, UT) and responses were collected between February 2018 and April 2020. The survey was advertised on a wide range of online platforms, including international academic mailing lists, staff and student mailing lists within Queen Mary University of London, as well as Twitter (https://www.twitter.com) and Reddit (https://www.reddit.com). Responses were obtained from 2,069 participants aged from 18 to 77 years (*M* = 23.6 years, *SD* = 8.8 years) and originating from a wide range of geographical areas (62 % North America, 25% Europe, 8% Asia, 3% Oceania, 1% Africa, 1% South America).

The resulting dataset required extensive manual cleaning comprising five main stages. First, we removed entries from participants who abandoned the survey study before providing any pieces of music, resulting in 1,398 entries being retained. Second, we removed entries that kept the default answers provided in the questionnaire. Third, we processed MEC onsets as follows: in the few cases where a participant indicated a time range rather than a specific time point, we took the start of the range as the onset of the MEC (e.g., 0:34–0:46 was replaced by 0:34); extraneous qualitative comments provided by a small number of participants about specific musical characteristics associated with MECs were removed (e.g., “0:26 where the guitar comes in” was replaced by 0:26); and a few entries in which the onset time provided was greater than the total track duration were removed. Fourth, we cleaned the URLs by removing additional qualitative comments and discarding non-valid URLs. Together, stages 2-4 retained 1,187 pieces of music associated with MECs, corresponding to 1,150 unique pieces with 2,028 MEC onsets. The fifth and final stage involved removing pieces that could not be accessed via the supplied URLs.

The final dataset consists of 1,806 MEC onsets in 1,019 unique pieces supplied by 402 individuals. The number of tracks reported by each participant ranged from 1 to 10 tracks (10 was the maximum permitted; median = 2, IQR = 2) while 988 tracks were reported by one participant only, 29 by 2 participants, and 2 by 3 participants. The number of MECs reported by each participant in each track ranged from 1 to 5 (5 was the maximum permitted; median = 1, IQR = 1). Combining MECs from different participants, the number of MECs per track ranged from 1 to 8 (median = 1, IQR = 1) while 611 tracks (60% of the total number) had exactly 1 report of MECs. Figure 4 provides descriptive statistics for the oChiM dataset including the most frequently occurring artists and genre tags; the distributions of track durations and MEC position within a track; and the distribution of vocal content and key distances (see Methods: Feature Extraction) across the dataset.

**Figure 4:**
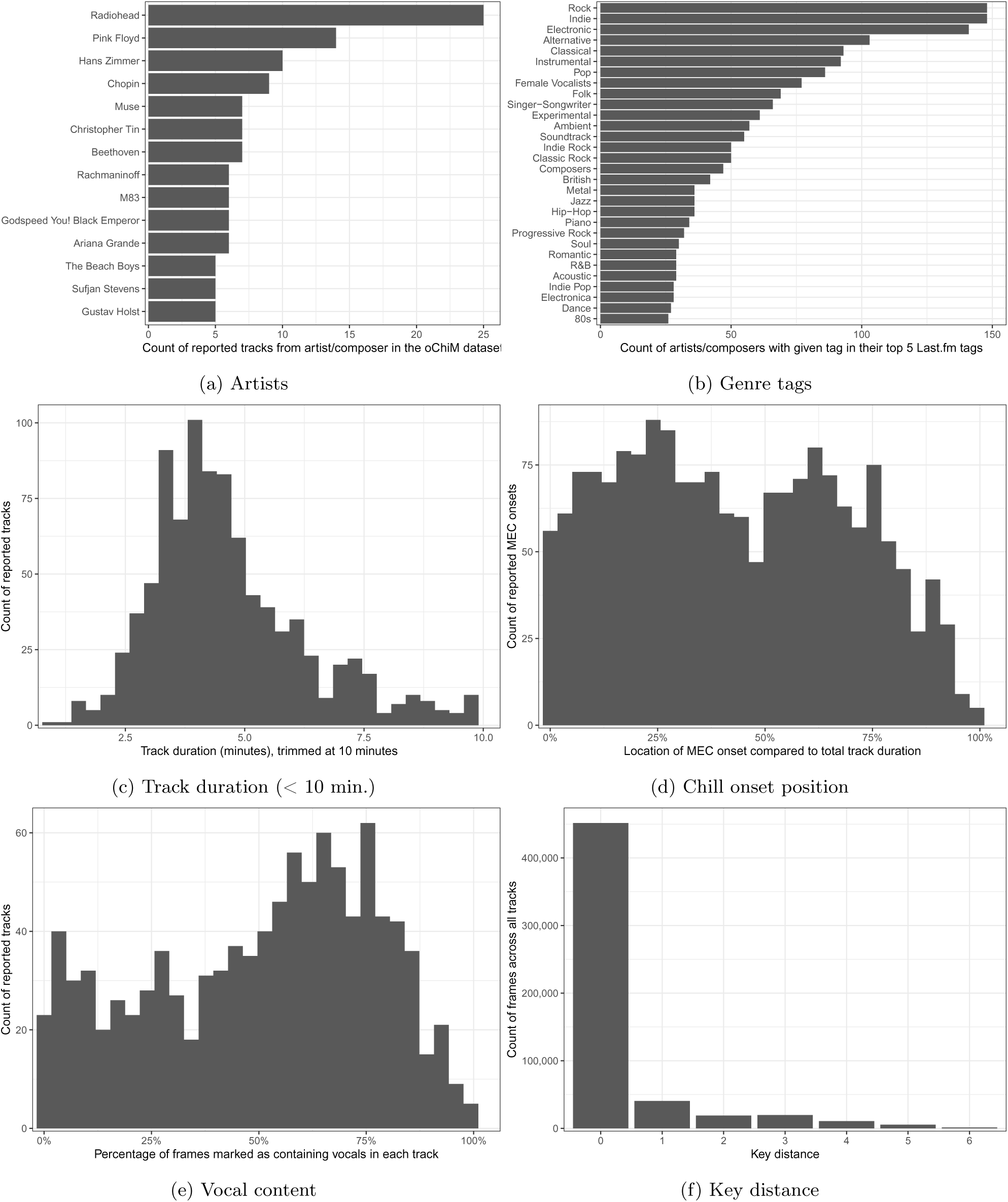
Descriptive statistics for the oChiM dataset. (a) Histogram of artists/composers appearing five or more times across the dataset. (b) Histogram of the 30 most frequently occurring tags acquired via the Last.fm API by searching for the artist name and extracting the top five user-submitted tags. (c) Histogram of track durations up to 10 minutes (1,596 tracks, 88% of the dataset); the remaining 210 tracks ranged up to 96 minutes with the majority (152 tracks) having a duration of 25 minutes or less. (d) Histogram of chill onset positions within a track, represented as percentage of track duration. (e) Histogram of vocal content, represented as percentage of frames marked as containing vocals in each track (see Methods: Feature Extraction). (f) Histogram of key distance across the dataset, representing distance on the cycle of fifths between estimated keys in successive 1 s windows (see Methods: Feature Extraction). Source data are provided as a Source Data file.

Overall the dataset consists primarily of Western popular and art music with a wide variety of styles represented including pop, rock, jazz, classical, electronic and folk. Note that because they are submitted by users these tags are likely to be noisy and incomplete. Track duration shows a peak at around 4 minutes but varies widely up to 96 minutes. Most tracks have some vocal content with very few in which no vocal content is detected at all but a reasonable minority containing less than 25% detected vocal content. Interestingly, the distribution of chill positions within tracks is bimodal. A speculative explanation would be that MECs early in the track reflect recognition of salient themes on first appearance whereas the second peak positioned around the golden ratio reflects the introduction of transitional or developmental material or perhaps the restatement of a theme following such material. Estimated key tends to be stable with changes of key being uncommon and those that do occur usually involving transitions to related keys.

### 4.2 Stimulus preparation

In preparation for auditory feature extraction, tracks were retrieved as WAV files from the URLs supplied by participants, downmixed to mono and downsampled to 44.1 kHz when necessary, using the *tuneR* R package [76]. RMS normalisation was applied to all tracks using the *soundgen* R package [77]: first, each track was linearly rescaled so that its peak amplitude was 0dB and the RMS amplitude of the track was computed; second, the track with the lowest RMS amplitude was found and every other track was linearly rescaled such that the RMS amplitude was equal to the lowest RMS amplitude.

### 4.3 Feature extraction

Based on the acoustic and musical elicitors reviewed in the Introduction, we extracted a range of features intended to simulate perception of loudness, roughness, brightness, spectral centroid and flux, event density, frequency range, crescendi, tonality, texture, expectation and presence of vocals. Due to the computationally intensive nature of extracting many features from many tracks, the scripts were run in parallel on multiple Linux servers at Queen Mary University of London. The whole feature extraction process is visualised in Figure 5, along with the source of the features and the elicitors they were intended to simulate.

**Figure 5:**
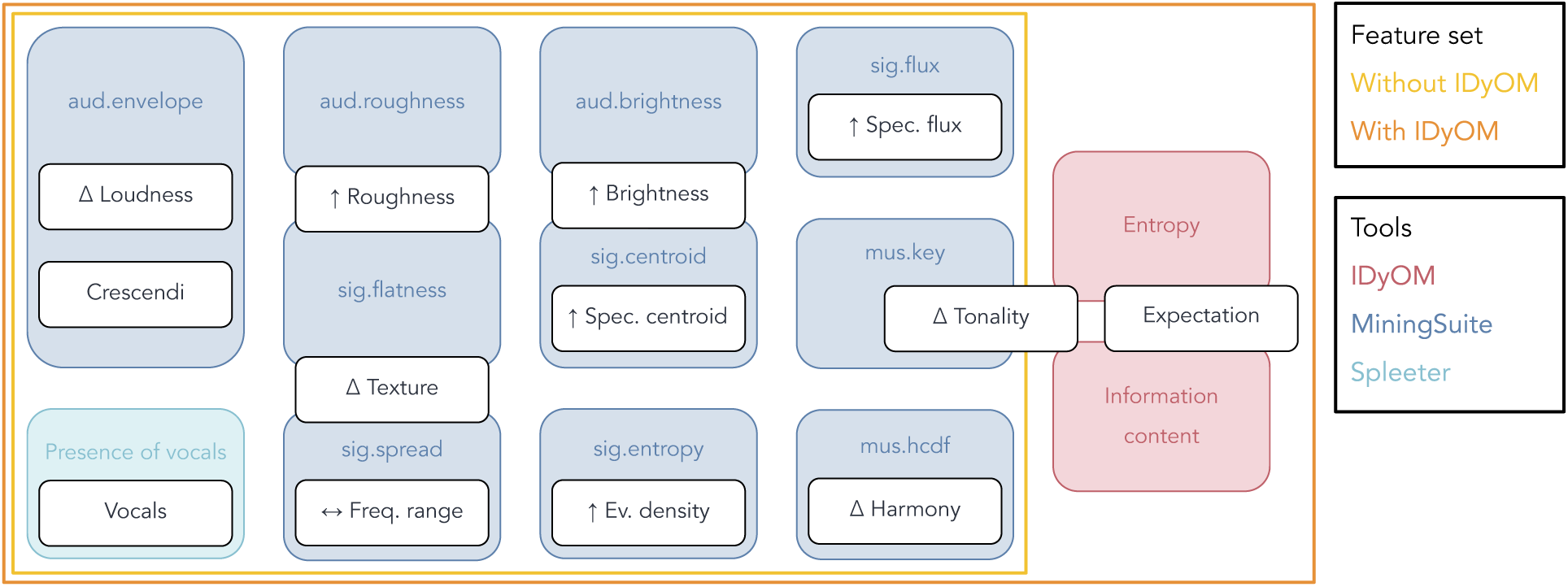
List of features extracted from each track. The features are depicted in boxes with coloured frames indicating the tool used to extract them. The acoustic and musical elicitors of MECs they aim to simulate are shown in white boxes (using the abbreviations *freq.* for “frequency”, *spec.* for “spectral”, and *ev.* for “event”), along with the hypothesised direction of their respective effects, as identified from prior research (see Introduction) and denoted by the following symbols: Δ for changes, *↑* for increases or elevated levels, and *↔* for expansions. The outer boxes represent the feature sets used for model training (see Supplementary Information for details). Source data are provided as a Source Data file.

#### 4.3.1 MiningSuite

Most features were extracted using *MiningSuite* [78], a MATLAB toolbox for the analysis of audio and music recordings, which expands on the methods provided by the commonly used *MIRtoolbox* [69]. Default settings for frame size were used as detailed below.

Loudness and crescendi were approximated with the *aud.envelope* function, which consists of a generic envelope extraction method, further processed following a model of human auditory perception [79]. The feature was extracted with a 10-ms frame size, corresponding to a 100-Hz sampling rate.

Spectral frame decomposition was performed using the *sig.spectrum* function, which applies a Fast Fourier Transform to the audio waveforms of each file, using a 50-ms sliding window with a 25-ms overlap. With the same sliding window size and overlap, we approximated roughness with the *aud.roughness* and *sig.flatness* features, respectively estimating sensory dissonance and spectral smoothness; brightness with the *aud.brightness* and *sig.centroid* features, the former capturing the amount of high-frequency energy in the signal and the latter more broadly capturing spectral centroid (the centre of mass of the spectrum–––another elicitor of interest); spectral flux with the *sig.flux* feature, calculating the spectral distance between successive frames; event density with the *sig.entropy* feature, computing the relative Shannon entropy of the input [80]; and frequency range with the *sig.spread* feature, capturing spectral variability.

Tonality was approximated with the *mus.key* feature, which estimates tonal centre by choosing the highest key candidate from a key strength curve, itself computed by correlating the chromagram of the signal with known key profiles [81, 82]. This feature was computed in a 1-s sliding window with a 0.5-s overlap. To approximate harmonic change, a six-dimensional tonal centroid was extracted using the *mus.tonalcentroid* function, corresponding to chord projections on the circle of fifths, minor thirds and major thirds, before being processed with the *mus.hcdf* harmonic-change detection function (HCDF), which returns the flux of the tonal centroid, using the default settings of a 743-ms sliding window with a 74.3-ms overlap.

#### 4.3.2 Spleeter

To generate a continuous, binary feature representing the presence of vocals, tracks were first processed using the *Spleeter* source separation Python library [83]. It provides pretrained models to perform source separation of a music track into two, four, or five stems containing separate instruments. For the present study, two-stem separation was conducted, resulting in two separate stems containing vocals and instrumental parts for each track.

We applied an amplitude threshold to the vocals stem. This produced a binary feature reflecting the absence or presence of vocals discarding the small amount of residual noise in the tracks. In practice, amplitude is characterised by a high degree of zero-crossing, which is not suitable for thresholding. Therefore, we computed the loudness of the vocals from their amplitude using the *soundgen* R package, which provides a function allowing the estimation of subjective loudness in sones (a psychoacoustic unit of perceived loudness) for each 20-ms sliding window with 50% overlap, resulting in a value capturing subjective loudness every 10 ms. To prevent the application of a loudness threshold from returning an overly sensitive vocals-detection feature, we applied a rolling maximum filter with a span of 510 ms for each track before finally applying a loudness threshold categorising vocals as present if above 2.5 sones. This thresholding process is visualised in Figure 6 and resulted in a continuous, binary feature representing the presence of vocals for each 10-ms frame.

**Figure 6:**
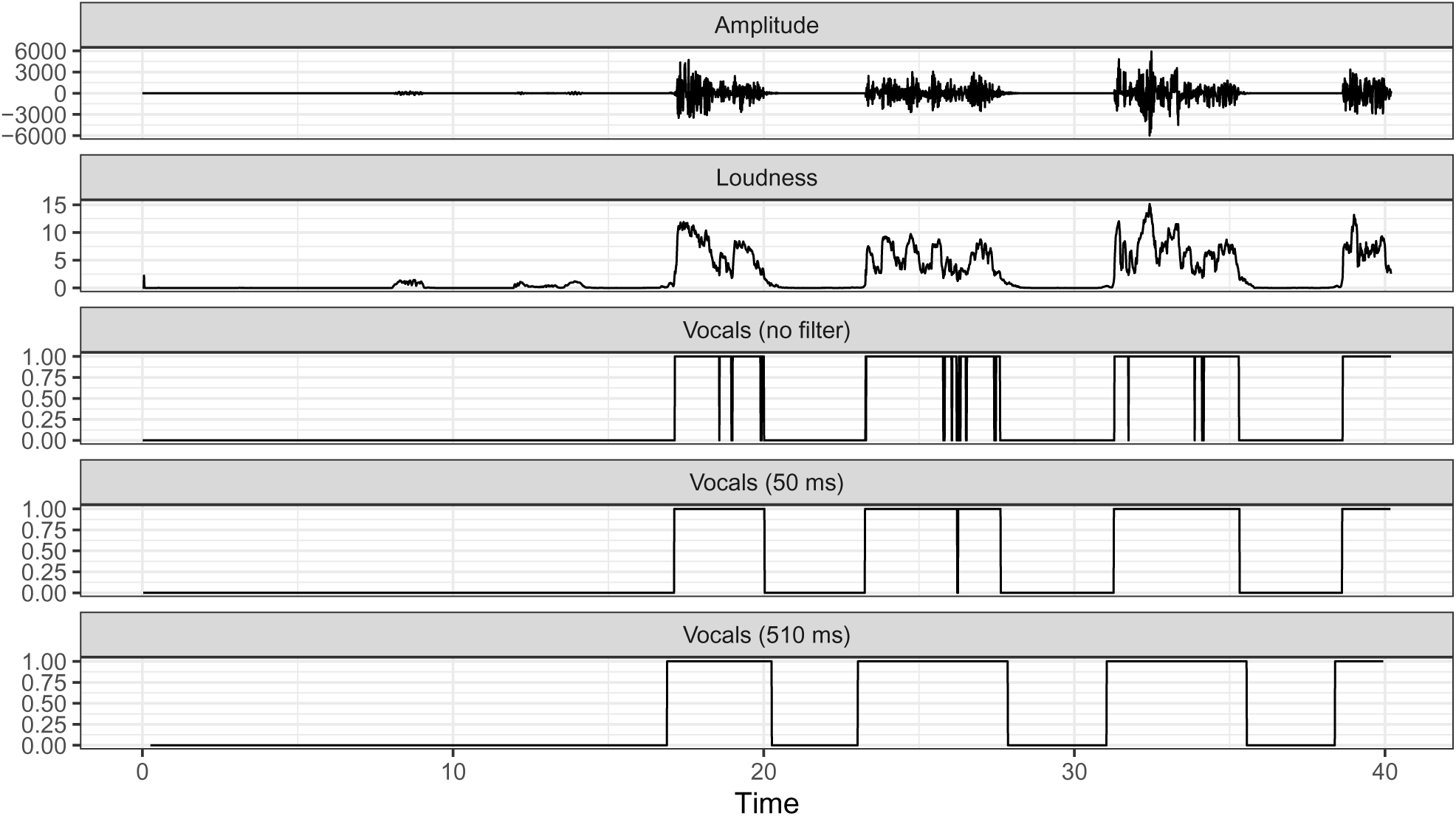
Vocals thresholding process. Data is shown for a short, 40-s excerpt to illustrate the thresholding process used to identify the presence of vocals. From top to bottom row are displayed 1) the amplitude of the waveform of the vocals stem extracted using Spleeter, 2) an estimate of subjective loudness, 3) the results of applying a 2.5-sones loudness threshold with no rolling maximum filter, 4) same as above with a rolling maximum filter with a 50-ms span, and 5) same as above with a 510-ms span. Several combinations of values were tested before the values used in the last plot were retained because they most closely approximated manual annotation in a representative set of examples. Source data are provided as a Source Data file.

Note that both the 510-ms value for the span of the maximum filter and the 2.5-sones value for the loudness threshold were chosen manually after comparing the outputs when altering these parameters. A systematic validation was not possible due to the absence of labelled data in our sample. However, we conducted several manual checks on a representative set of recordings and deemed the feature resulting from these parameters as close as possible to what would have resulted from manual annotation of the tracks. Importantly, while this feature is sensitive to differences in loudness between tracks, such concerns were mitigated by the fact that RMS normalisation was conducted prior to this step in the analysis.

#### 4.3.3 MELODIA

To simulate high-level melodic expectation, it was necessary to extract melodies in MIDI format from each audio track. First, the time series of continuous frequency values in Hz was extracted for each melody using the *MELODIA* plugin [84] for *Vamp* (https://vamp-plugins.org) via its associated *vamp* Python library (https://pypi.org/project/vamp) MELODIA enables the estimation of the fundamental frequency of the pitch of the primary melody in polyphonic tracks, and was therefore particularly well suited for this task.

MELODIA has been shown to perform well on multiple melody extraction benchmarks covering a range of musical styles (pop, rock, jazz, classical, north Indian vocal music and Chinese karaoke) achieving a general accuracy of 0.75 [84] measured as the proportion of frames for which either the ground truth and prediction were both unvoiced or the predicted F0 was within 50 cents of the ground truth F0 [85]. A simple heuristic provided by the author of MELODIA (https://github.com/justinsalamon/audio_to_midi_melodia) was reimplemented in R to quantise pitch frequencies into discrete MIDI notes: converting each value in Hz to its closest MIDI note before applying a median filter with a 250-ms span to remove some of the underlying data noise, and finally discarding notes shorter than 100 ms. The resulting MIDI notes were stored in a text file suitable for the next step of the feature extraction process, while note onsets were stored in a reference file separately to convert the extracted features back to time series suitable for model training.

#### 4.3.4 IDyOM

The Information Dynamics of Music (IDyOM) model [44, 57], was used to generate information-theoretic measures of melodic expectation from the sequences of MIDI events extracted by MELODIA. IDyOM generates probabilistic predictions for musical events using variable-order Markov models, which learn from statistical regularities in symbolic musical sequences such as MIDI representations. Two outputs are relevant here: *entropy*, a measure of uncertainty about which event is predicted to come next given the current context; and *information content*, the negative log probability, a measure of the surprisal of the event. For instance, if at time *t −* 1 in a melody, IDyOM is very certain about which note comes next, this will be reflected by low entropy for the note occurring at time *t*. If the actual next note is indeed very predictable, this will be reflected by low information content, but if it is unpredictable, information content will be high. IDyOM can be configured to learn incrementally from a given piece of music (via its short-term model, capturing dynamic effects on expectation), to generate predictions based on prior learning from a large corpus of music (the long-term model, capturing schematic effects) or both. IDyOM provides a multiple-viewpoint system, which allows sequences to be predicted using a range of different representations of the musical input including, for example, abstract representations of pitch interval and scale degree for predicting the pitch of a note. IDyOM has been the subject of substantial empirical evaluation and has proved capable of simulating expectation, complexity, arousal and pleasure in music perception across a range of experiments [47, 49, 50, 58].

In the present study, the *cpint* viewpoint (chromatic pitch interval) was derived from the *cpitch* viewpoint (MIDI pitch number) to assess the entropy and information content of each MIDI event. The model used a combination of the short-term and long-term models, with the long-term model being trained on a set of folk ballads from Nova Scotia, Bach chorale melodies, and German folk songs from the Essen Folk Song Collection, consistent with the geographical origins of most participants in the MEC onset survey [for a description of this corpus, see 44, Table 3.2]. Training on this corpus has been shown in previous research to produce a model that simulates perceptual expectations of Western listeners [44, 47, 49, 57–59]. The resulting entropy and information content values associated with each MIDI event for each track were linked back to the note onsets, allowing these values to be converted to continuous features synchronised with the tracks.

### 4.4 Feature preparation

#### 4.4.1 Key distance

Most features required additional processing to capture the hypothesised elicitors motivated by prior research. Tonality, notably, is only thought to affect the occurrence of MECs in occasional cases of changes in tonality. However, the current feature extracted with *mus.key* only captured the tonal centre at a given time, which should have no bearing on the occurrence of MECs. Therefore, we computed key distance from this feature, following the hypothesis that changes in tonality would be more conducive to experiencing MECs than tonality itself. To do so, we first smoothed the feature by applying majority voting with a 3.5-s span, before assigning a value to tonality changes based on distance in the circle of fifths. For instance, if the tonal centre at time *t −* 1 was C, the key distance at time *t* would be 0 if the tonal centre remained C, 2 if it changed to D, or a maximum of 6 if it changed to F#. The feature was then upsampled to 100 Hz for consistency with the other features, replacing the newly introduced missing values with their nearest existing key distance value.

#### 4.4.2 Interpolation and smoothing

Cubic spline interpolation was applied to all the other auditory features extracted with MiningSuite to match this 100-Hz frame rate, except for envelope, already sampled at 100 Hz. Interpolation was only allowed to fill the gaps between two existing values, to prevent the method from returning wildly unlikely values at the beginning and end of each track. Following this, all features (including envelope) were smoothed using a 50-ms median filter.

#### 4.4.3 First-order difference

As discussed earlier and shown in Figure 5, many MEC elicitors refer to changes, not specific values, in acoustic and musical properties. To reflect this, first-order differences were computed for each feature and included in the analyses such that if a feature had a value of 3 at time *t −* 1 and a value of 7 at time *t*, its first-order difference at time *t* would be 7 *−* 3 = 4.

#### 4.4.4 Segmentation

Reducing the quantity of frames being considered for analysis was necessary to speed up computations. Tracks were segmented into larger frames, over which summary statistics were computed to investigate central tendencies and variability for each feature. Segmentation into 200-ms and 500-ms frames was compared with 500-ms performing best (see Supplementary Information) and retained for the main analysis. Mean and standard deviation were computed for each frame, resulting in four dimensions for each original feature: *µ*_0_ and *σ*_0_ (mean and standard deviation of the original values) and *µ*_1_ and *σ*_1_ (mean and standard deviation of the first-order difference). The way key distance was computed prevented us from calculating standard deviations as these would not contain meaningful information. In the present study, *µ*_0_ can be thought of as the original feature, *σ*_0_ as the variation in that feature, *µ*_1_ as the rate of change in the feature, and *σ*_1_ as the variation in that rate of change. For conciseness, we refer to *σ*_0_ and *σ*_1_ as measures of variability.

We predicted that, for envelope, HCDF and key distance, changes on a slower time scale might better capture their hypothesised role as elicitors of MECs. Summary statistics for these features were therefore also computed using a 2-s sliding window with 50% overlap, and upsampled to the target frame size (200-ms or 500-ms) for inclusion in the analyses. The full list of features is shown in Table 1, and the computed features are available in the oChiM dataset accompanying the present study.

### 4.5 Analysis

#### 4.5.1 Permutation tests

To identify patterns in the behaviour of each feature around the onset of MECs, we ran a series of permutation tests [for a similar approach, see 20]. Permutation tests provide a convenient non-parametric way of constructing a test statistic, comparing the observed data to a null distribution generated by permuting the samples. Here, we examined the impact of each feature on MECs by evaluating how unlikely their values were around the onset of MECs, when compared to other moments within the tracks that were not reported as producing MECs.

To achieve this, all 20-s excerpts centred around the MEC onsets from each track were extracted. We only retained complete excerpts, meaning that MEC onsets within the first or last 10 s of each track were discarded. Rather than choosing individual control excerpts for each MEC excerpt, every track was split into sequential 20-s excerpts and we discarded any excerpt that was incomplete or overlapping with an MEC excerpt, retaining all remaining excerpts as controls. This process resulted in two unbalanced sets of 20-s excerpts, capturing 10 s before and after each MEC onset for the MEC set and most other moments within the tracks for the control set. The test statistic was computed for each frame of each feature, consisting of the difference between the average values for excerpts causing MECs and control excerpts. Two-tailed permutation tests were run by randomly permuting the excerpts while keeping the same number of excerpts in each set, using Monte Carlo estimation with 5,000 replications. We only ran the permutation tests on the data with a 500-ms frame size to limit the number of comparisons, and we used Bonferroni correction within each feature to further mitigate the fact that, even with a 500-ms frame size, 41 significance tests would be required for each feature.

#### 4.5.2 Principal component analysis

Before training classifiers to examine whether or not MEC onsets could be predicted using acoustic and expectation-based predictors, it was important to assess collinearity between the predictors. Figure 7 shows a correlation matrix for all the predictors which highlights strong collinearities between some pairs of features. It is not surprising that positive correlations exist between key distance, HCDF and envelope and the corresponding variants computed over a larger sliding window (key distance+, HCDF+ and envelope+). However, there are also strong positive collinearities between several of the acoustic measures of spectral components of timbre (brightness, flatness, roughness, as well as spectral centroid, flux, entropy and spread). Furthermore, some of these spectral measures (roughness and spectral flux) correlate positively with envelope as a measure of loudness suggesting that louder parts of the music also show greater dissonance and spectrotemporal variation. Finally, it is worth noting that the expectation-based measures (melodic IC and entropy) do not correlate either with each other or with any of the acoustic measures.

**Figure 7:**
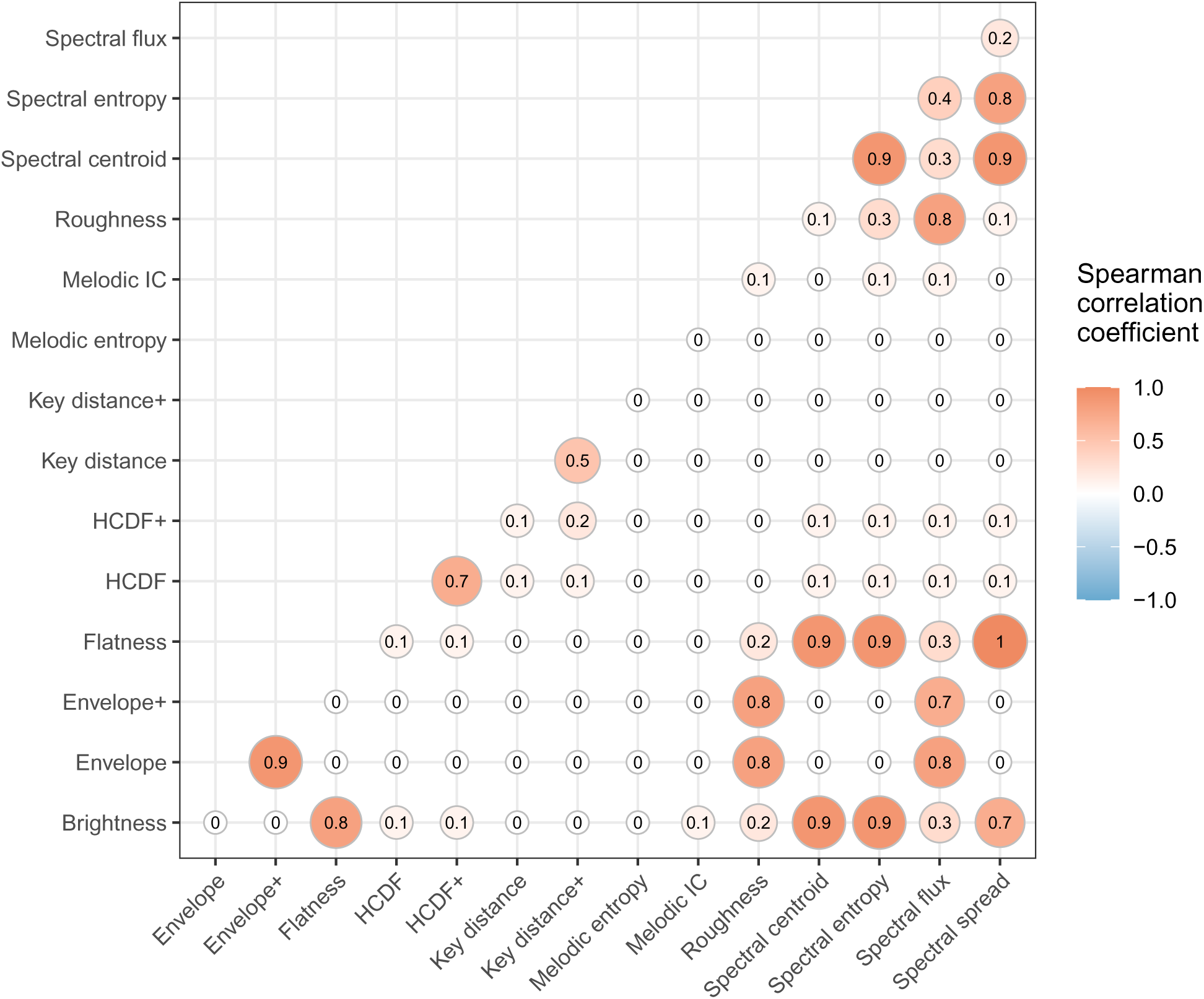
Spearman correlation coefficients between feature values for 500 ms frames in every track in the oChiM dataset for each simulated acoustic and expectation-based elicitor of MECs. Source data are provided as a Source Data file.

Given this collinearity between the features, we ran a principal component analysis (PCA), projecting each data point into a new space, the dimensions of which are known as principal components. PCA enables dimensionality reduction when retaining the smallest number of components that account for as much variance in the data as possible. The results of the PCA are discussed here for simplicity, as they are not crucial to the interpretation of the results reported. We retained 15 principal components, accounting together for 78% of the cumulative proportion of variance explained for by the PCA. We did not attempt to interpret how the features were grouped together into specific principal components, due to the difficulty of assigning meaning to such complex combinations of weights without introducing unnecessary subjectivity. However, we stored feature loadings on each principal component and proportion of variance explained by each principal component for later analyses of feature importance.

#### 4.5.3 Support vector machine

Classification analyses were run using the support vector machine (SVM). For classification tasks using an SVM, model training consists of finding the hyperplane that maximises the distance between the two classes and is used as a decision boundary to make predictions. SVMs are widely used for classification tasks and were chosen in the present analysis because they are relatively easy to interpret and have the added benefit of being more forgiving in terms of statistical assumptions than logistic regression–––another commonly used classification method. The main drawback is that the SVM does not consider the sequential nature of the data. Therefore, the SVM was compared with a GMM-HMM classifier based on a hidden Markov model (HMM) predicting continuous time series modelled as Gaussian mixtures (a Gaussian mixture model or GMM), which is sensitive to temporal patterns in data; the results showed that the SVM outperforms the GMM-HMM (see Supplementary Information), so we restrict ourselves to the SVM in the main analyses.

SVMs were trained using the *scikit-learn* Python library [86] with a linear kernel, no random feature selection, stopping criterion set at 10*^−^*^5^ and by adjusting weights in inverse proportion to class frequencies to account for class imbalance. To further mitigate class imbalance and the imprecision in the time resolution of reported MEC onsets, we assigned positive labels to all frames within 1 s around each MEC onset (i.e., a 2-s excerpt centred on each MEC), corresponding to the resolution with which participants were asked to report MECs in the survey. For instance, with a 500-ms frame size, instead of considering a single frame for an MEC onset at *t* = 40 s, we considered frames at *t* = 39, 39.5, 40, 40.5, and 41 s as positive labels for the classification task. SVMs were trained to distinguish positive (MEC onsets) from negative (control) labels and the regularisation parameter for linear SVMs (using a logarithmic scale ranging from 10*^−^*^24^ to 10^4^, with even-numbered exponents only) was optimised using manual grid search. We used five-fold cross-validation, using for each fold 60% of the tracks as a training set, 20% as a validation set to pick the best-performing model for testing, and 20% as a testing set to return final performance metrics for the learning parameters which performed best across all five folds. Overall, given 1,806 MECs, this resulted in 8,840 positive samples, averaging 1,768 per fold and 5,304 per training set (each consisting of 3 folds).

Performance was assessed by comparing ground truth and predictions in a fixed time grid, splitting each track into consecutive 5-s segments and marking these segments as active if they included at least one frame representing a predicted or actual MEC onset, and inactive otherwise. A given 5-s segment was categorised as a true positive if it was active in both the ground truth and the SVM predictions. This resulted in the evaluation of 82,310 segments (2,634 positive and 79,676 negative in the ground truth). To evaluate classification performance, we computed precision, recall and the F-measure — the harmonic mean of precision and recall — as well as balanced accuracy, true and false positive rates, and the area under the ROC curve (AUC). In a typical classification task, the AUC is calculated by modifying the classification threshold. However, linear SVMs do not return the probabilities associated with the predictions for each frame by default. To obtain these probabilities, we applied Platt scaling [87] on the decision function and computed the AUC with *scikit-learn* by adjusting the classification threshold over the percentiles of the returned probabilities. After observing the results, we also decided to compute F*_β_*, which applies an additional weight *β* to the F-measure in order to disproportionately favour precision or recall over the other. We picked a value of 2 for *β*—a standard value to signify that we considered recall twice as important as precision in this classification task to avoid optimising for small differences in precision.

The results reported in the Supplementary Information show that the best performing model was an SVM with 500-ms frame size and high-level expectation-based features included.

#### 4.5.4 Feature importance

The final step of the analysis was to compute feature importance. To simplify the interpretation of results, feature importance was only extracted from the best-performing model across all combinations of frame size, feature set and model type (see Supplementary Information). Since the best performing model was an SVM, the method described below is specific to feature importance extraction from the parameters of a trained SVM.

Using Platt scaling involves a second layer of cross-validation within each cross-validation fold. We refer to the folds from this second layer as *SVM folds* to differentiate them from the cross-validation folds used in the workflow detailed above. First, feature coefficients were extracted from the parameters of the trained models. Because we were interested in assessing feature importance, as opposed to trying to interpret the directionality of the coefficients (which was more relevant to the permutation test analyses), absolute values were taken, therefore preventing coefficients from cancelling each other out when averaged over SVM folds, if directionally different. These absolute values for the coefficients of each feature were averaged over each SVM fold, resulting in a set of five positive averaged coefficients for each feature–––i.e., one averaged coefficient per cross-validation fold.

However, in this case, the features were principal components obtained with PCA. To map the coefficients back to the original features, we applied weights based on the proportion of variance explained and feature loadings on each principal component, thereby assigning a share of the influence of each principal component on the models to its constituent features based on how much variance was accounted for by the principal component. To do so, we simply took the feature loadings on each principal component, multiplied by the proportion of the variance explained by that principal component and multiplied by the coefficient extracted from the models for that principal component. We then summed the coefficients for each feature across all principal components, resulting, again, in a single coefficient per feature for each of the cross-validation folds (but this time, for each of the original features, as opposed to principal components).

The resulting coefficients were normalised to ensure that they were comparable. Since the magnitude of the coefficients differed between cross-validation folds but the folds were of equal size, we rescaled the coefficients linearly for each fold such that 0 corresponded to the least contributing feature for that fold and 1 to the most contributing feature. Finally, we averaged these values across the five cross-validation folds, resulting in a single feature importance value between 0 and 1 for each original feature used for PCA before model training. These values do not exactly correspond to a rank but rather to a continuous spectrum of feature importance, where 0 would correspond to the feature with the lowest feature importance on all five cross-validation folds, and 1 to that with the highest feature importance.

## Data availability

The Onsets of Chills in Music (oChiM) dataset generated in this study has been deposited in the OSF database under accession code doi:10.17605/osf.io/x59fm [https://doi.org/10.17605/osf.io/x59fm]. Source data are provided with this paper.

## Acknowledgements

This research was supported by the EPSRC and AHRC Centre for Doctoral Training in Media and Arts Technology [EP/L01632X/1].

## Author contributions

CRediT: Conceptualization (RdF, MTP); Data curation (RdF); Formal analysis (RdF); Funding acquisition (MTP, EB); Investigation (RdF); Methodology (RdF, MTP, EB, AC); Project administration (RdF, MTP); Resources (RdF); Software (RdF); Supervision (MTP, EB); Validation (RdF); Visualization (RdF); Writing – original draft (RdF, MTP); Writing – review & editing (EB, AC).

## Inclusion and ethics

Ethics approval for this research was granted by the Ethics of Research Committee at Queen Mary University of London (QMREC1752). All participants gave informed consent before taking part.

