## Supplementary Information for "Melodic expectation as an elicitor of music-evoked chills"

#### Introduction

The supplementary information contains further details of the MEC classification analysis.

For the classification, all analyses were run with two segmentation parameters: once for segmentation into 200-ms frames and once for 500-ms frames, because there was no information to determine a priori which frame size would work best to investigate the occurrence of MECs. In addition to these two segmentation parameters, two feature sets were evaluated. The first set did not include the IDyOM features simulating high-level expectation while the second set did. This was done to evaluate the effect of accounting for expectation in the predictive performance of the models. For clarity, we hereafter refer to these differences in model training as differences in *frame size* (200-ms or 500-ms), *feature set* (without or with IDyOM features) and *model type* (described below).

Therefore, four PCAs were conducted with centring and scaling—one for each combination of frame size (200 ms or 500 ms) and feature set (with or without IDyOM). We retained 12 principal components with eigenvalues above one for each PCA using the first feature set (without IDyOM) and 15 principal components for each PCA using the second feature set (with IDyOM), both regardless of frame size. Finally, we compared two kinds of classifier: one based on a hidden Markov model and the other based on a support vector machine, which are covered below.

#### Hidden Markov models

As opposed to Markov chains, which model the probabilities of sequences of observable states, hidden Markov models (HMMs) model the probabilities of hidden states, which themselves drive the probability distributions of observable events. HMMs are specified by a set of hidden states, the transition probabilities between these states, an initial probability distribution for these states, a set of observations and the observation likelihoods associated with each state, also called emission probabilities—the probability that an observation was generated by a specific state [for excellent introductions to HMMs, see 1, 2].

In its most simple form, for a univariate sequence of observations, an HMM associates each hidden state with a specific emission distribution (e.g., normal distribution with given mean and standard deviation) of the observed values in that sequence. In practice, we are often faced with multivariate sequences of observations, and instead of using multivariate Gaussian emissions, which might be limited in how accurately they can represent observations, Gaussian mixture model emissions (GMMs) are preferred. GMM-HMMs are particularly well suited to modelling auditory events due to their flexibility and sequential nature, and have successfully been applied to speech recognition [see 3] and to segmentation, genre classification, sequence prediction and event detection in music [e.g., 4, 5].

In the present case, the observations correspond to the multidimensional sequence of principal components, of which different configurations were modelled by GMMs, each governed by a different hidden state of the HMM. It was assumed that specific sequences of hidden states gave rise to the occurrence of MECs—as opposed to a single hidden state representing MECs specifically.

HMMs are characterised by three problems [1, 2]: the likelihood problem (determining the likelihood of a specific sequence of observations given the HMM and observations), the decoding problem (discovering the best hidden state sequence given the HMM and observations—not relevant to the present study because we did not attempt to interpret the hidden states themselves), and the learning problem (learning the HMM parameters given the states and observations). The present automatic MEC onset detection task consisted of two steps. First, we trained two separate HMMs (learning problem), one trained on excerpts centred around the MEC onsets, and the other on control excerpts. Second, we presented these trained HMMs with new excerpts to obtain the likelihood of each excerpt according to each HMM (likelihood problem). If the HMM trained on excerpts causing MECs returned the highest likelihood, the excerpt was categorised as inducing MECs and vice versa. Implementation details are provided below.

As with the permutation tests, this approach required extracting two sets of excerpts, because HMMs are best trained on short sequences of equal length. The exact same procedure was used, selecting excerpts centred around the MEC onsets, and control excerpts sequentially through the rest of each track. However, instead of 20-s excerpts, we tested two excerpt durations: 2 s (with a 200-ms frame size only), and 5 s (with 200-s or 500-s frame sizes), to speed up computations while ensuring a reasonable amount of frames for each excerpt. *Excerpt size* (2 s or 5 s) thus corresponds to the last aspect of model training we controlled, in addition to frame size, feature set and model type.

Survey participants reported MEC onsets with a 1-s resolution. We consequently decided to augment the training data by also extracting excerpts categorised as causing MECs for each frame within 1 s of the original MEC onset. For instance, with a 500-ms frame size, instead of getting a single excerpt for a MEC onset at $t$ = 40 s, we extracted five excerpts at $t$ = 39, 39.5, 40, 40.5, and 41 s. This also allowed us to reduce the class imbalance, with control excerpts outnumbering excerpts causing MECs by a ratio of 100:1.

Using the *pomegranate* Python library [6], two GMM-HMMs (MECs and control) were trained for each combination of feature set (with or without IDyOM), frame size (200 ms or 500 ms), and excerpt size (2 s or 5 s), using manual grid search by iterating over the number of hidden states and the number of Gaussian mixtures with which the models should be trained. More specifically, to save time on this computationally intensive training process, we trained each model using odd numbers of states and mixtures (ranging from 1 to 17) before testing the even numbers of states and mixtures nearest to the best-performing model. This process is illustrated in Figure 1. We used five-fold cross-validation, using for each fold 60% of the tracks as a training set, 20% as a validation set to pick the best-performing model for testing, and 20% as a testing set to return final performance metrics for the combination of learning parameters that performed best across all five folds. As with feature extraction, model training was run in parallel on university-provided servers.

The process for training an individual GMM-HMM was as follows. All data were concatenated ignoring their sequential nature to identify a cluster for each hidden state using k-means and initialise the parameters of the corresponding GMM using such a cluster. The model was then initialised with a uniform probability transition matrix before training began using the Baum-Welch algorithm [see 1]. Regularisation was applied by setting a transition and emission pseudocount of 0.1 and an edge and distribution inertia of 0.1 [see 6]. If the model failed to converge, generally due to underflow errors, training was attempted once more. If unsuccessful, training was abandoned and a new model was trained for the next step of the grid search (as in Figure 1).

Full tracks, split into excerpts centred around each consecutive frame (as opposed to excerpts split sequentially for model training), were used for model validation. For each excerpt from each track in the validation set, a log probability value was obtained for each pair of trained GMM-HMMs (one trained on excerpts causing MECs and the other trained on control excerpts), using the forward algorithm [see 1]. If the GMM-HMM trained on excerpts causing MECs returned the highest log probability for the tested excerpt, that excerpt was predicted as an occurrence of MEC, and vice versa. For each fold, this validation process resulted in a univariate time series of binary frame-wise predictions for each track. As with the SVM classifiers described in the Methods section of the main paper, evaluation was conducted on 5-s consecutive segments marked as active if they included at least one frame representing an MEC onset, and inactive otherwise.

In a typical classification task, the AUC is calculated by modifying the classification threshold. In the present analysis, however, predictions were not based on a classification threshold but resulted from comparing log probabilities between two GMM-HMMs. To emulate the principle of classification threshold, we collected all frame-wise log-probability differences and extracted their percentiles. Each percentile was used as a proxy for a threshold by generating a new set of predictions based on whether or not the difference between the log probabilities of both GMM-HMMs was higher or lower than that percentile. This allowed us to collect 100 pairs of true positive and false positive rates (one for each percentile), which were used to compute the AUC with the *scikit-learn* Python library [7]. For the validation sets, only AUC was used to select the combination of learning parameters that performed best across all five folds. For each fold, the GMM-HMMs trained using these learning parameters were then used on the testing set to compute the AUC. All other performance metrics were computed for the classification threshold (i.e., the log probability difference threshold) that returned the highest $F_\beta$-value. These metrics were then averaged across all five folds to return the final model-performance metrics.

### Support vector machines

The SVM classifiers are described in the Methods section of the main paper.

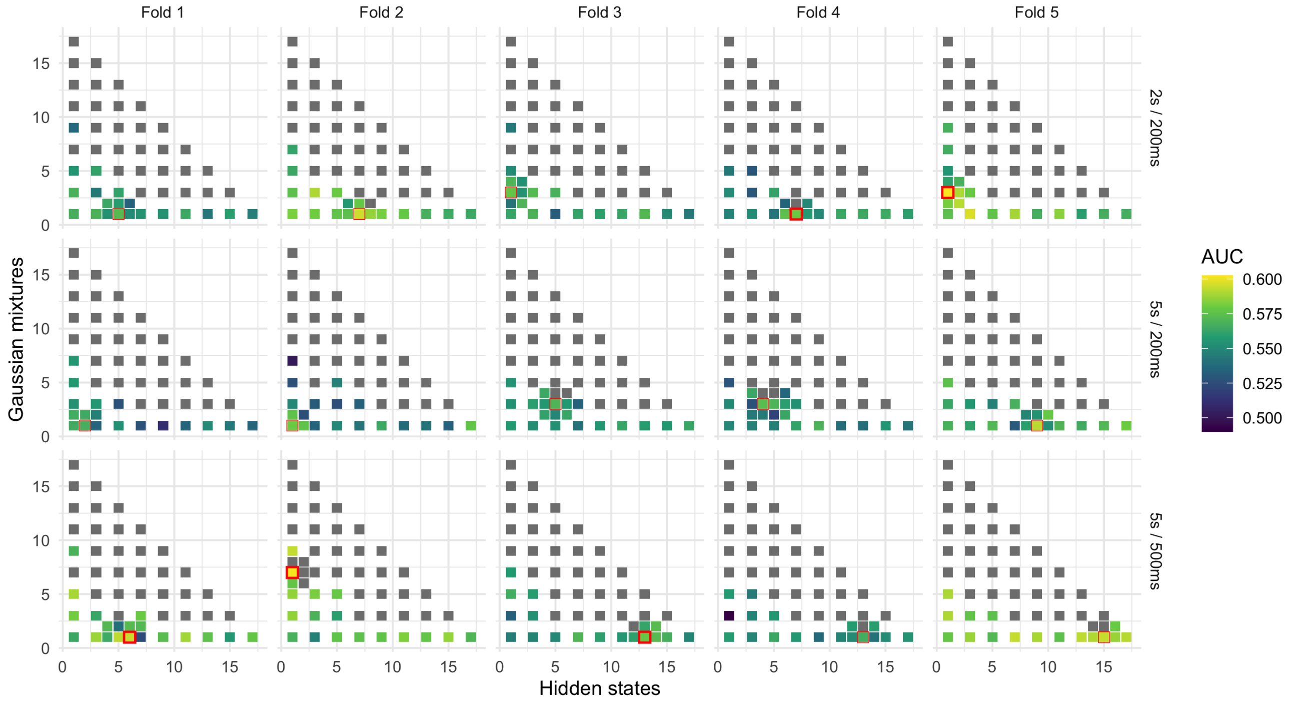


Figure 1: Grid search for GMM-HMM training. In this example, GMM-HMMs were trained on the feature set including IDyOM features. Two GMM-HMMs (MECs and control) were trained for each combination of the number of hidden states (bottom axis), Gaussian mixtures (left axis), cross-validation fold (top axis) and excerpt size and frame size (right axis). Model performance was evaluated as the area under the receiver-operating characteristic curve (AUC). Odd numbers of states and mixtures were first used for training. For each combination of fold, excerpt size and frame size, additional models were trained using the even numbers of states and mixtures closest to the best-performing models (highlighted in red), sometimes resulting in enhanced performance (e.g., Fold 1, 5-s excerpt size, 500-ms frame size). Grey cells signal models which failed to converge. The best-performing GMM-HMMs for each fold are highlighted with a thicker, red border. Source data are provided as a Source Data file.

| **Model** | **IDyOM** | **Frame size** | **Excerpt size** | **Evaluation metrics** | | | | | |
|---|---|---|---|---|---|---|---|---|---|
| | | | | AUC | $F_\beta$ | $F$ | $P$ | $R$ | $BA$ |
| GMM-HMM | ✗ | 200 ms | 2 s | 0.568 | 0.156 | 0.072 | 0.038 | 0.692 | 0.555 |
| | | | 5 s | 0.579 | 0.161 | 0.075 | 0.040 | 0.682 | 0.564 |
| | | 500 ms | 5 s | 0.560 | 0.159 | 0.073 | 0.038 | **0.800** | 0.561 |
| | ✓ | 200 ms | 2 s | 0.562 | 0.157 | 0.071 | 0.037 | 0.788 | 0.556 |
| | | | 5 s | 0.564 | 0.156 | 0.071 | 0.037 | 0.782 | 0.552 |
| | | 500 ms | 5 s | 0.574 | 0.164 | 0.076 | 0.040 | 0.755 | 0.572 |
| SVM | ✗ | 200 ms | - | 0.591 | 0.166 | 0.077 | 0.041 | 0.716 | 0.578 |
| | | 500 ms | - | 0.592 | 0.166 | 0.078 | 0.041 | 0.713 | 0.579 |
| | ✓ | 200 ms | - | 0.594 | 0.166 | 0.077 | 0.041 | 0.724 | 0.579 |
| | | 500 ms | - | **0.597** | **0.167** | **0.078** | **0.041** | 0.693 | **0.580** |

Table 1: Evaluation metrics for the SVM and GMM-HMM classifiers. Best performance for each metric (column) is shown in bold. AUC = Area under the receiver-operating characteristic curve, $F_\beta$ = weighted F-measure, $F$ = F-measure, $P$ = Precision, $R$ = Recall, $BA$ = Balanced accuracy.

### Results

The SVM and GMM-HMM were compared with 200-ms and 500-ms frame sizes, with and without expectation-based features (IDyOM). For performance evaluation, we computed precision, recall and the F-measure—the harmonic mean of precision and recall as well as $F_\beta$ with a value of 2 for $\beta$ (see Methods). In addition, we computed balanced accuracy as well as true positive and false positive rates to extract the AUC. Table 1 shows a comparison of the SVMs and GMM-HMMs with and without IDyOM, with 200-ms and 500-ms frame sizes, The best-performing classifier was an SVM using a 500-ms frame size and including IDyOM. This was used for the main analysis.

A final analysis assessed the benefit of combining four classes of statistic for each feature: the mean ($\mu$) and variability ($\sigma$) of the zeroth (0) and first derivative (1) (see Methods). The mean feature importance shown in Figure 3 fell from $\mu_0$ (0.597) $>$ $\sigma_1$ (0.522) $>$ $\sigma_0$ $>$ (0.485) $>$ $\mu_1$ (0.213) and a Kruskal-Wallis test showed a significant effect of class, $\chi^2(3) = 21.95, p < .001, \eta^2 = 0.327$. Running the optimal SVM analysis only with the best performing class (mean feature values: $\mu_0$) resulted in poorer overall performance (AUC: 0.559; $F_\beta$: 0.163; F: 0.074; Balanced accuracy: 0.554) albeit with elevated recall but lower precision (Precision: 0.039; Recall: 0.826). One reason for this is that different features vary in the relative importance of each class of statistic. For example, referring to Figure 3, mean level ($\mu_0$) is more important for spectral flatness, centroid and spread while variability ($\sigma_0$ and $\sigma_1$) is more important for roughness and envelope.
